# Haplotypes of common SNPs can explain missing heritability of complex diseases

**DOI:** 10.1101/022418

**Authors:** Gaurav Bhatia, Alexander Gusev, Po-Ru Loh, Bjarni J. Vilhjálmsson, Stephan Ripke, Schizophrenia Working Group of the Psychiatric Genomics Consortium, Shaun Purcell, Eli Stahl, Mark Daly, Teresa R de Candia, Kenneth S. Kendler, Michael C O’Donovan, Sang Hong Lee, Naomi R. Wray, Benjamin M Neale, Matthew C. Keller, Noah A. Zaitlen, Bogdan Pasaniuc, Jian Yang, Alkes L. Price

**Author notes:** Correspondence should be addressed to G.B. or A.L.P.

## Abstract

While genome-wide significant associations generally explain only a small proportion of the narrow-sense heritability of complex disease (*h*^2^), recent work has shown that more heritability is explained by all genotyped SNPs (*h*_*g*_^2^). However, much of the heritability is still missing (*h*_*g*_^2^ < *h*^2^). For example, for schizophrenia, *h^2^* is estimated at 0.7-0.8 but *h*_*g*_^2^ is estimated at ∼0.3. Efforts at increasing coverage through accurately imputed variants have yielded only small increases in the heritability explained, and poorly imputed variants can lead to assay artifacts for case-control traits. We propose to estimate the heritability explained by a set of haplotype variants (haploSNPs) constructed directly from the study sample (*h*_*hap*_^2^). Our method constructs a set of haplotypes from phased genotypes by extending shared haplotypes subject to the 4-gamete test. In a large schizophrenia data set (PGC2-SCZ), haploSNPs with MAF > 0.1% explained substantially more phenotypic variance (*h*_*hap*_^2^ = 0.64 (S.E. 0.084)) than genotyped SNPs alone (*h*_*g*_^2^ = 0.32 (S.E. 0.029)). These estimates were based on cross-cohort comparisons, ensuring that cohort-specific assay artifacts did not contribute to our estimates. In a large multiple sclerosis data set (WTCCC2-MS), we observed an even larger difference between *h*_*hap*_^2^ and *h*_*g*_^2^, though data from other cohorts will be required to validate this result. Overall, our results suggest that haplotypes of common SNPs can explain a large fraction of missing heritability of complex disease, shedding light on genetic architecture and informing disease mapping strategies.

## Introduction

Genome-wide association studies (GWAS) have been extremely successful in identifying robust associations between single nucleotide polymorphisms (SNPs) and complex traits, providing important biological insights. However, despite the large number of loci with genome-wide significant associations, these loci explain only a small proportion of the heritability of complex traits^1^. For example, for schizophrenia the heritability estimated from related individuals (*h^2^*) is 0.7-0.8^2^, and the heritability explained by all loci with genome-wide significant associations (*h*_GWAS_^2^) to schizophrenia is only 0.03^3^.

The gap between the heritability estimated from related individuals^4^ (*h^2^*), and that explained by genome-wide significant associations (*h*_GWAS_^2^)—termed the “missing heritability^5,6^”—is partially explained by causal variants that have not achieved genome-wide significance in GWAS of current sample sizes^7^. Indeed, the heritability of schizophrenia explained by all genotyped SNPs (*h*_*g*_^2^) has been estimated at ∼0.3^8^ (all values on liability scale^9^). While this is substantially larger than *h*_GWAS_^2^ of 0.03, it leaves much of *h^2^* unaccounted for. This pattern of *h*_GWAS_^2^< *h*_*g*_ ^2^ < *h^2^* is observed across a broad set of complex traits^1^. Thus, despite a clear polygenic signal, a substantial portion of the heritability remains missing. One possible explanation for this missing heritability is inflation in *h^2^* estimates due to genetic interactions^10^ or shared environment^11,12^. A second possibility is that currently unobserved rare genetic variants explain a significant portion of the variance of studied traits^7,13^.

A logical first step in including untyped genetic variants in analyses would be to perform imputation using a higher coverage reference panel^14^. Increasing coverage through imputation is an essential part of association and fine-mapping. While accurately imputed SNPs do not typically explain more heritability than genotyped SNPs alone^8,15^, including low-accuracy imputed SNPs can explain substantially more of the heritability of quantitative traits (see Discussion). However, it is unclear whether poorly imputed SNPs can be included in analyses of case-control traits, due to the severe danger of assay artifacts inflating the resulting estimates^9^. Additionally, imputation will not include the contributions of variants that are absent or rare in reference panels. This is particularly important for large-effect rare variants that will be at dramatically higher frequency in ascertained case-control studies than in the population.

To better understand the contributions of rare and low frequency variants (MAF < 0.05), we estimated the heritability of complex traits explained by haplotype variants (haploSNPs) constructed from within the sample. This is based on the idea that two individuals who are identical by state for a long haplotype of DNA are more likely to also share unobserved rare variants linked to the haplotype^16^. This allows us to tag rare variants whose phenotypic effects are poorly captured by genotyped or accurately imputed SNPs.

Using simulations involving real genotypes, we demonstrated that haploSNPs can explain substantially more heritability than SNPs alone, primarily due to superior tagging of unobserved rare causal variants. We applied our method to >35,000 schizophrenia cases and controls from the Psychiatric Genomics Consortium 2 (PGC2-SCZ) and obtain an estimate of *h*_hap_^2^ = 0.64 (S.E. 0.084), representing a substantial increase over *h*_*g*_^2^ = 0.32 (S.E. 0.029). These estimates were based on cross-cohort comparisons, ensuring that cohort-specific assay artifacts did not contribute to our estimates. We separately applied our method to >14,000 multiple sclerosis cases and controls from the Wellcome Trust Case Control Consortium 2 (WTCCC2-MS). We observed an even larger difference between *h*_*hap*_^2^ and *h*_*g*_^2^, though data from other cohorts will be required to validate this result.

## Results

### Overview of methods

To assess whether rare variants are contributing to the gap between the heritability explained by common SNPs (*h*_*g*_^2^) and the total narrow-sense heritability (*h^2^*), we estimated the heritability explained by haplotype variants (haploSNPs) constructed from multiple SNPs (*h*_*hap*_^2^). haploSNPs are haplotypes of adjacent SNPs excluding a subset of masked sites that arise from skipped mismatches (see below). Individuals are considered to carry 0,1, or 2 copies of the haploSNP if none, one or both of their chromosomes matches the haplotype at all unmasked sites.

Using computationally phased genotypes^17^, we build haploSNPs from all pairs of phased chromosomes in the sample, independently of phenotype. We start with a SNP at which the two chromosomes match, and extend one SNP at a time. The haploSNP is extended until a terminating mismatch—a mismatch that cannot be explained as a mutation on a shared background. Terminating mismatches are detected as violations of the 4-gamete test between the haploSNP being extended and the mismatch SNP. We also limit each haploSNP to a maximum length of 50kb (see Online Methods), which our simulations suggest captures much of the rare variant tagging while ensuring a relatively small number of haploSNPs (avoiding large S.E. in *h_hap_^2^*). We have released open-source software implementing the method for building haploSNPs (see Web Resources).

Once all pairs of chromosomes are analyzed, the set of haploSNPs is treated as a new set of variants that can be analyzed using standard techniques. We analyzed these haploSNPs in a linear mixed-model framework^7^ (see Online Methods). We used PCGC regression^18^ (a generalization of Haseman-Elston regression^19^) instead of REML to estimate the components of heritability explained by this specific set of haplotype variants, avoiding the bias of REML methods (e.g. GCTA) under extreme case-control ascertainment^18,20^ (see Online Methods). We stratified by MAF^21,22^ to avoid bias due to MAF-dependent genetic architectures, employing a MAF-stratified PCGC regression approach. We employed an efficient implementation of PCGC regression, which we have released as an open-source software package (see Web Resources). For our analyses where multiple cohorts of similar ancestry were available (i.e. our schizophrenia analysis), we estimated heritability using comparisons of individuals from different cohorts to avoid biases due to cohort-specific assay artifacts.

### Simulations using real genotypes

We first sought to assess the degree to which haploSNPs improved tagging of rare causal variants using a large sample of sequenced individuals from the UK10K project (see Web Resources). Starting with 3,565 unrelated individuals sequenced at 235k SNPs on chromosome 22, we masked all SNPs that were not present on the Illumina Human660-Quad chip, and considered the remaining 6,477 SNPs as “genotyped SNPs” in our analysis. We computationally phased these genotyped SNPs, and built a set of 937k haploSNPs using our approach. To compare the tagging efficiency of well-imputed SNPs and haploSNPs, we used Impute2^23^ to impute 24k SNPs with INFO > 0.9 from a 1000 Genomes reference panel^24^ (see Online Methods). For each SNP in the sequence data, we compared the best tag from the set of genotyped SNPs, well-imputed SNPs, or the set of haploSNPs (see Online Methods). We observed much higher tagging efficiency for haploSNPs (average best tag from haploSNPs of *r*^2^ = 0.54) compared with well-imputed or genotyped SNPs (average best tags of *r*^2^ = 0.34, 0.28). This improvement was much larger than would be expected by chance given the large number of haploSNPs (see Online Methods) and was driven primarily by improved tagging for sequence SNPs with MAF < 0.05 (see Figure 1).

**Figure 1.**
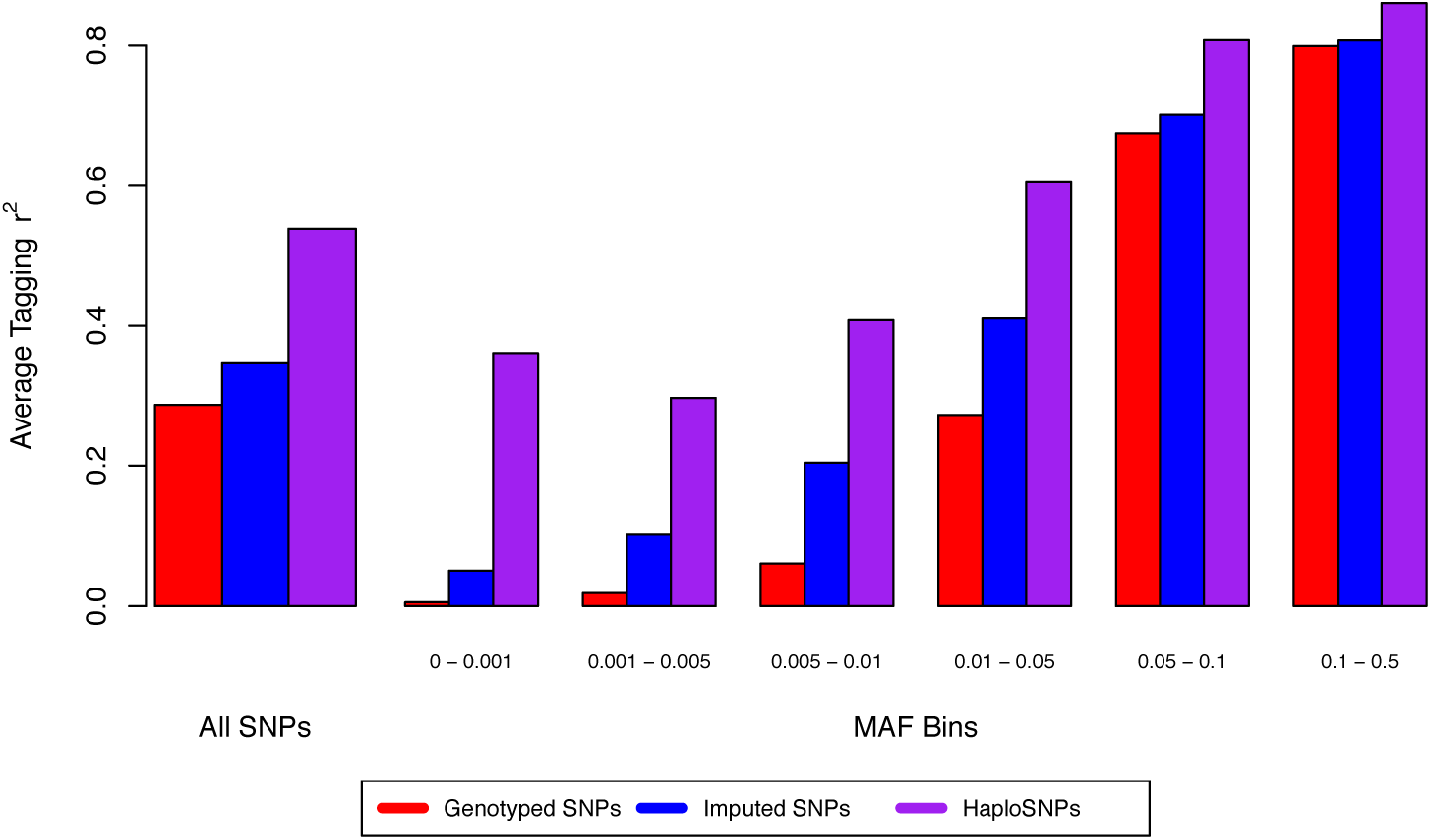
Assessing the degree of improved tagging of underlying sequence variants when using haploSNPs as compared with genotyped SNPs alone. The largest magnitude of increase is for SNPs with MAF < 0.05, consistent with our observations of increased heritability.

We next sought to ensure that our estimation procedure produced robust estimates of the heritability explained by haploSNPs (*h*_*hap*_^2^). Using the set of haploSNPs from chromosome 22 generated above, we simulated quantitative traits by selecting a subset of the haploSNPs as causal variants (see Online Methods). We simulated multiple genetic architectures, each with causal variants chosen from a different subset of the haploSNP MAF distribution (i.e. MAF 0-0.05) (total *h^2^* = 0.5 for all simulations). These simulations allowed us to verify that heritability estimates from haploSNPs closely matched the heritability used to simulate phenotypes (Figure 2; see Online Methods).

**Figure 2.**
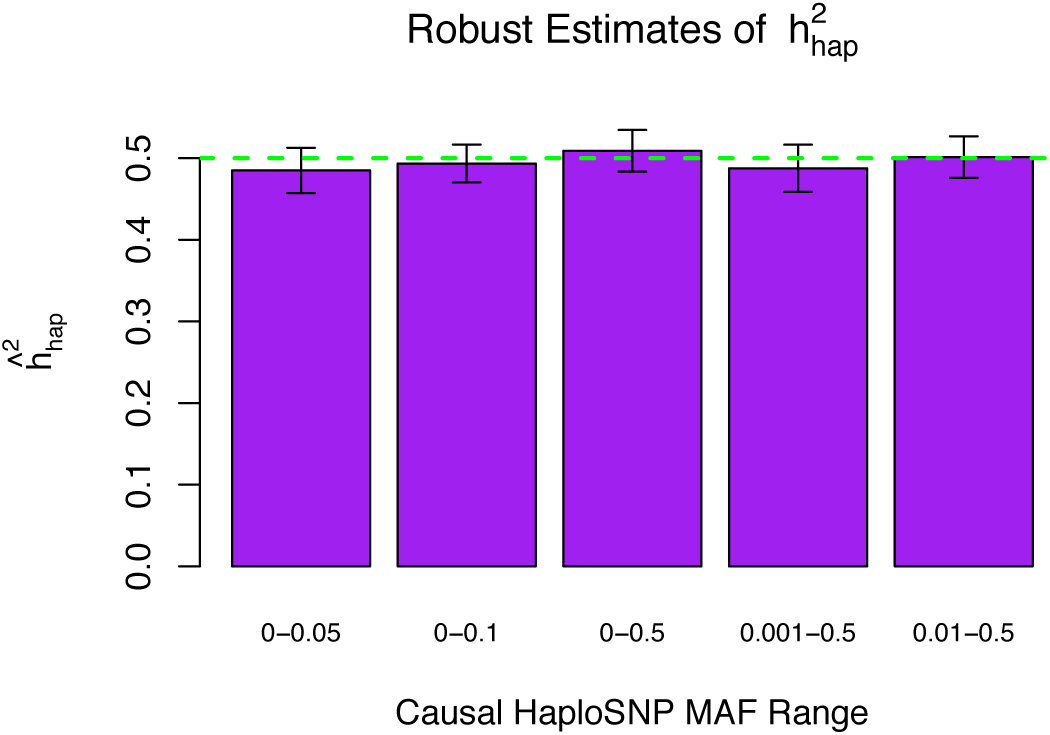
Estimating the heritability explained by haploSNPs for phenotypes simulated from causal haploSNPs. The true heritability in all simulations is 0.5. Because we knew the true heritability, we could evaluate whether the resulting estimates reproduce the expected value. Each of the estimates is averaged over 100 simulated phenotypes. The error bars indicate the 95% confidence interval surrounding this mean value (computed as twice the standard error of the mean).

To assess our ability to explain additional heritability using haploSNPs, we simulated phenotypes from UK10K sequence SNPs (instead of haploSNPs as above) using multiple genetic architectures, each with causal variants chosen from a different subset of the MAF distribution (total *h^2^* = 0.5 for all simulations). We compared the heritability explained by genotyped SNPs and haploSNPs (see Figure 3). We note that well-imputed SNPs did not explain additional heritability and were subject to known LD-bias^15,21^ (see Figure S1). Our results show that genotyped SNPs explain the bulk of heritability when causal variation is common: for example, if causal SNPs are drawn with MAF > 0.01, then *h*_*g*_^2^ = 0.36 (S.E. 0.005) and *h*_*hap*_^*2*^ = 0.39 (S.E. 0.015). However, as the contribution of rare variants to phenotype increases, haploSNPs can explain substantially more heritability than SNPs alone. For example, if causal SNPs are drawn with MAF < 0.05 then *h*_*g*_^2^ = 0.038 (S.E. 0.002) and *h*_*hap*_^*2*^ = 0.24 (S.E. 0.016). We explored the effect of altering the maximum length threshold on resulting heritability estimates (see Figure S2); based on these results, we chose to use haploSNPs with MAF > 0.1% and length shorter than 50kb to maximize tagging while limiting the number of haploSNPs generated (and thus the S.E. on *h*_*hap*_^*2*^). Results on explaining additional heritability using haploSNPs were similar using GCTA^25^ (see Figure S3), as expected in the absence of case-control ascertainment. We note that available sample sizes of sequence data did not allow for simulation of strong case-control ascertainment (e.g. prevalence < 0.01) with realistic haplotype/rare-variant structure.

**Figure 3.**
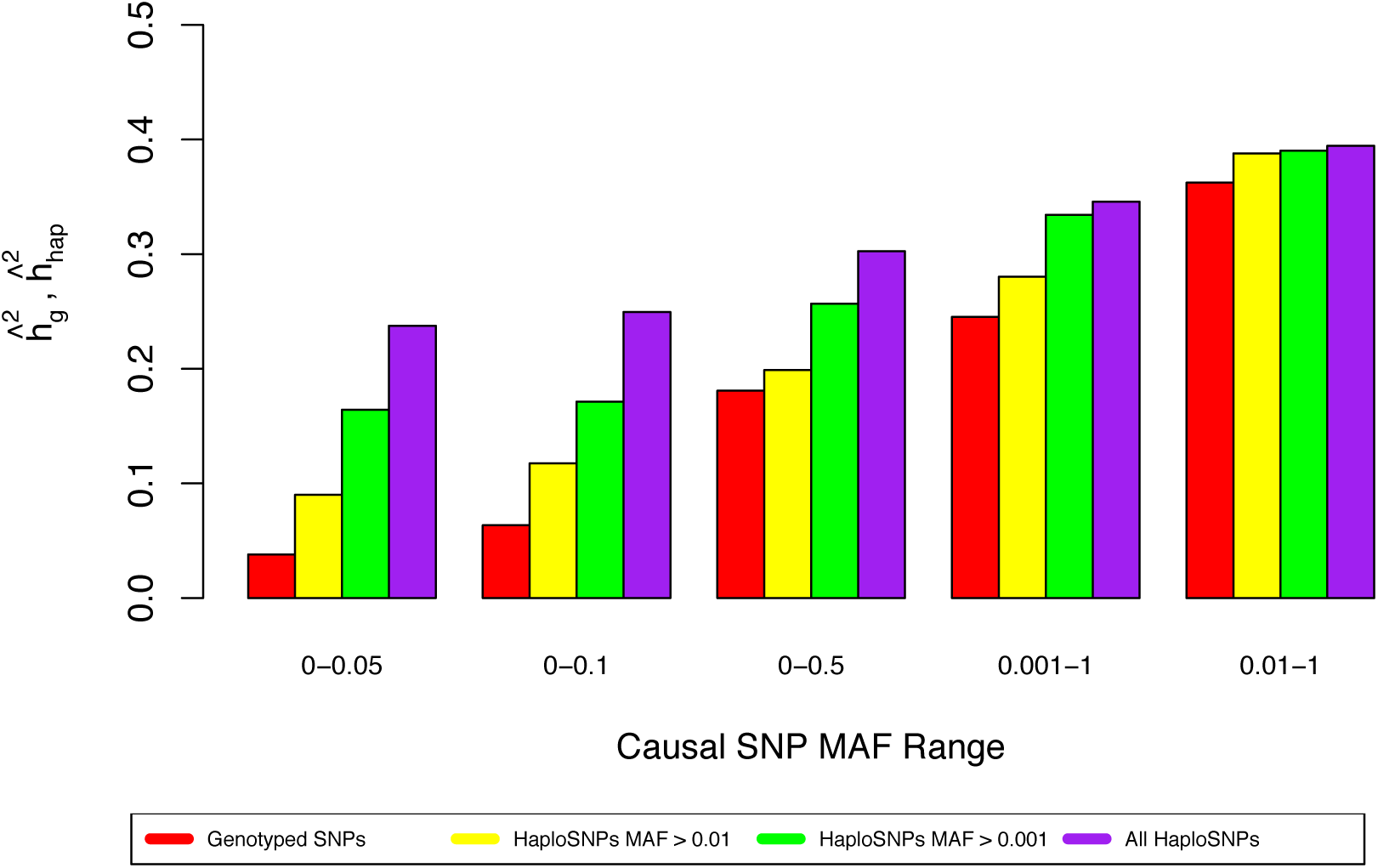
We compared the performance of haploSNPs of less than 50kb (at a variety of MAF thresholds) to that of SNPs alone. For each causal MAF range, 100 phenotypes were simulated. Ranges were (in order from left) MAF < 0.05, 0.1, and 0.5 and MAF > 0.001 and 0.01. All phenotypes have a total heritability of 0.5. Averages across simulations are reported in the figure. We note that these ranges are overlapping so that a causal variant with MAF < 0.05, may also be causal in the simulation with MAF < 0.01. It is clear that SNPs will tag the bulk of the heritability in common variant dominated genetic architectures. However, for rare variant dominated architectures, haploSNPs can provide a substantial increase in the heritability explained.

### Analysis of schizophrenia

We analyzed the genome-wide heritability explained by genotyped SNPs and haploSNPs in PGC2-SCZ data^3^. (We note that well-imputed SNPs have been shown to not explain substantially more heritability than genotyped SNPs alone in previous analyses of PGC-SCZ data^8^.) We first meta-analyzed estimates for each of ten cohorts of European ancestry with >1,000 individuals, for a total of >35,000 individuals (this is distinct from the cross-cohort analysis described below). In each cohort, we applied stringent quality control to genotyped SNPs, obtaining an average of 461k genotyped SNPs in each cohort (see Online Methods and Table 1). We computationally phased these genotypes^17^ and constructed an average of 36.3M haploSNPs in each cohort. We stratified haploSNPs according to MAF, and estimated the heritability explained jointly with SNPs using MAF-stratified PCGC regression^18^ with 7 variance components (see Online Methods) using a disease prevalence of 1%^26^. For two studies of treatment resistant schizophrenia a disease prevalence of 0.3% was used. (We note that the choice of disease prevalence affects the absolute estimates of *h*_*g*_^2^ and *h*_*hap*_^2^, but not their relative values.) To counter effects of population stratification, we included 140 principal components in all analyses—20 from each of 6 haploSNP MAF ranges and 20 from the SNPs (see Online Methods). Heritability estimates for these cohorts were meta-analyzed using inverse variance weighted averaging producing estimates of *h*_*g*_^2^ = 0.29 (S.E. 0.020), consistent with previous reports^26^, and *h*_*hap*_^*2*^ = 0.72 (S.E. 0.071) for haploSNPs with MAF > 0.001 (all estimates on the liability scale) (see Table 1). We note that the larger standard error for *h*_*hap*_^*2*^ is expected due to the large number of haploSNPs. Estimates of *h*_*hap*_^*2*^ from GCTA were slightly lower than those from PCGC regression (see Table S1), potentially reflecting the known downward bias of REML estimates due to case-control ascertainment^18^.

**Table 1.** Summary of cohort-specific results in Schizophrenia

| Cohort ID | $h_{\text{hap}}^2$ | S.E. | $h_g^2$ | S.E. | N | SNPs | HaploSNPs |
| --- | --- | --- | --- | --- | --- | --- | --- |
| ajsz | 0.36 | 0.32 | 0.26 | 0.10 | 2,229 | 600k | 21.8M |
| boco | 1.46 | 0.28 | 0.30 | 0.07 | 3,847 | 387k | 26.2M |
| clm2 | 0.80 | 0.11 | 0.32 | 0.03 | 7,649 | 346k | 29.1M |
| clo3 | 1.16 | 0.32 | 0.37 | 0.07 | 3,950 | 487k | 39.5M |
| gras | 1.75 | 0.54 | 0.28 | 0.11 | 1,740 | 321k | 17.1M |
| irwt | 1.91 | 0.40 | 0.34 | 0.08 | 2,200 | 589k | 26M |
| mgs2 | 0.59 | 0.18 | 0.26 | 0.05 | 4,605 | 478k | 28.9M |
| s234 | -0.47 | 0.31 | 0.04 | 0.08 | 3,314 | 441k | 16.4M |
| swe5 | 0.54 | 0.21 | 0.33 | 0.06 | 3,866 | 477k | 33.3M |
| swe6 | -0.14 | 0.33 | 0.16 | 0.11 | 1,838 | 483k | 24.5M |
| **Meta-analysis** | **0.72** | **0.07** | **0.29** | **0.02** | **35,238\*** | **461k\*\*** | **36.3M\*\*** |

\*The number of individuals is summed across cohorts.
\*\*The number of SNPs and haploSNPs are averaged across cohorts.
We list data for each cohort that is included in our meta-analysis. We used 10 cohorts with $N > 1,000$ . The meta-analyzed estimate of $h_{\text{hap}}^2$ is significantly larger than the estimate of $h_g^2$ , though assay artifacts that differentiate cases and controls are a concern with these estimates. We note that negative estimates are expected due to statistical noise and that constraining estimates to the plausible 0-1 range would introduce bias in the meta-analysis; however, no estimates were statistically significantly less than 0.

Given the large increase in heritability explained (i.e. between *h*_*g*_^2^ and *h*_*hap*_^*2*^) we were concerned that assay artifacts differentially affecting cases and controls within each cohort^9,27^ could be inflating our estimates. To investigate this possibility, we performed a cross-cohort heritability analysis^8,28^ in two subsets of cohorts of homogenous ancestry. The first subset was of Swedish ancestry and consisted of five cohorts with a total of 2,819 schizophrenia cases and 2,911 controls. The second subset was of British ancestry and consisted of two cohorts with a total of 5,406 Treatment resistant schizophrenia cases and 5,284 controls (see Table S2). To maximize sample size, the Swedish subset included some cohorts with N < 1,000 that were not included in our analysis above. Following stringent QC, we analyzed genotypes at 310k and 303k SNPs in each subset, respectively (see Online Methods). To enable cross-cohort analyses within each subset, haploSNPs were rebuilt; 21.5 and 26.8 million haploSNPs were generated in the Swedish and British subset, respectively. Within each subset, we estimated heritability using MAF-stratified PCGC regression based only on pairs of individuals from different cohorts, instead of all pairs of samples (see Online Methods). Thus, unless assay artifacts are shared across multiple cohorts, they would not impact our heritability estimates. Meta-analyzed cross-cohort heritability estimates for the Swedish and British subsets were *h*_*g*_^2^ = 0.32 (S.E. 0.029) and *h*_*hap*_^*2*^ = 0.64 (S.E. 0.084), with the bulk of the increase coming from haploSNPs with MAF < 0.05 (see Table 2). These estimates are only slightly smaller than our overall estimates reported above, though we cannot rule out a small amount of inflation in those estimates due to assay artifact. To be conservative, we report the cross-cohort estimates as our final estimates. We note that the inclusion of rare haploSNPs does not decrease the contribution of common SNPs to phenotypic variance (see Table S3), suggesting key roles for both common and rare genetic variation in the genetic architecture of schizophrenia.

**Table 2.** Summary of cross-cohort results in Schizophrenia

| Model Components | Meta-Analyzed |  | Swedish |  | British |  |
| --- | --- | --- | --- | --- | --- | --- |
| | $h_{\text{hap}}^2$ | S.E. | $h_{\text{hap}}^2$ | S.E. | $h_{\text{hap}}^2$ | S.E. |
| haplo > 0.001 + SNPs | 0.64 | 0.084 | 0.36 | 0.146 | 0.77 | 0.103 |
| haplo > 0.005 + SNPs | 0.48 | 0.063 | 0.35 | 0.102 | 0.57 | 0.080 |
| haplo > 0.01 + SNPs | 0.42 | 0.044 | 0.29 | 0.077 | 0.49 | 0.054 |
| haplo > 0.05 + SNPs | 0.32 | 0.029 | 0.28 | 0.044 | 0.35 | 0.038 |
| SNPs Only* | 0.32 | 0.029 | 0.27 | 0.041 | 0.35 | 0.037 |

\*This is an estimate of $h_g^2$
To address the possibility that assay artifacts are impacting our estimates of $h_{\text{hap}}^2$ we estimated the heritability explained by haplotypes across cohorts with the same ancestry.
We considered one Swedish subset, and one British subset and meta-analyzed estimates across these cohorts.

To assess whether haploSNPs contribute to known genome-wide significant loci, we analyzed 99 loci containing genome-wide significant associations^3^ after merging adjacent loci (see Online Methods). The heritability explained by both SNPs and haploSNPs in these loci was estimated at *h_hap_^2^* = 0.042 (S.E. 0.018), which is not significantly larger than the heritability explained by SNPs alone (*h*_*g*_^2^ = 0.037; S.E. 0.007); these estimates were based on a cross-cohort analysis as described above. These results are consistent with prior reports^15^ that the causal allele frequency spectrum at GWAS loci is skewed towards common variants. We separately assessed whether haploSNPs might serve as better tags for underlying causal variants by evaluating whether the best haploSNP *χ*^2^ statistic was significantly larger than the best SNP *χ*^2^ statistic within the published locus. We observed an average increase in *χ*^2^statistics of 3.5 and 3.0 in the Swedish and British subsets (mean % increase: 55% and 38%), respectively. After correcting for the larger number of haploSNPs tested (see Online Methods), we found that this increase was statistically significant (K-S test P=3.9 × 10^−4^). This suggests that, despite not explaining additional heritability, haploSNPs do more effectively tag causal variants at associated loci. This distinction is explained by the fact that a linear combination of SNPs in the locus may tag a haploSNP well (so that heritability explained by the haploSNP is included in estimates of *h*_*g*_^2^) even though individual SNPs in the locus may not tag the haploSNP well.

### Analysis of multiple sclerosis

We analyzed the genome-wide heritability explained by genotyped SNPs and haploSNPs in the WTCCC2-MS data set^20,29^. We used extremely stringent quality-control filters to avoid inflation of genome-wide estimates by large numbers of small artifactual effects^9,15^. Because we could not validate our results across cohorts, the QC thresholds used for this data-set were more stringent than those used in our analysis of PGC2-SCZ (see Online Methods). Following QC, this data set consisted of 14,526 individuals (9,315 cases + 5,211 controls) genotyped at 349k SNPs (see Online Methods). We computationally phased this set of genotyped SNPs^17^ and built a set of 36.3 million haploSNPs. We note that there is a large ancestry mismatch between WTCCC2 MS cases and controls as a consequence of the set of samples that are publicly available^20^. While we also performed ancestry-matched analyses (see below), for our primary analysis we analyzed all individuals and included 140 PC covariates—20 from each of 6 haploSNP MAF ranges and 20 from the SNPs in our analyses (see Online Methods). We estimated *h*_*g*_^2^ = 0.20 (S.E. 0.019) and *h_hap_^2^* = 0.67 (S.E. 0.063) for haploSNPs with MAF > 0.001 (all estimates on the liability scale) (see Table 3). These estimates were produced using MAF-stratified PCGC regression^18^ with 7 variance components using a disease prevalence of 0.1%. Estimates of the heritability explained by haploSNPs using MAF-stratified GCTA^7,25^ were subject to known REML bias and varied widely as a function of ascertainment of cases^18,20^ (see Table S4). As expected, well-imputed SNPs from the 1000 Genomes Project^24^ did not explain substantially more heritability than genotyped SNPs (non-significant increase of 0.01; see Supplementary Note). While inclusion of SNPs with low imputation accuracy can explain more heritability of quantitative traits (see Discussion), it is unclear whether this approach is applicable to case-control traits due to the severe danger of inflation due to assay artifact^9^.

**Table 3.** Summary of results in Multiple Sclerosis

| **Multiple Sclerosis** |  |  |  |  |
| --- | --- | --- | --- | --- |
|  | **All (N=14,526)** |  | **Ancestry Matched (N=8,039)** |  |
| **Model Components** | **$h_{\text{hap}}^2$** | **S.E.** | **$h_{\text{hap}}^2$** | **S.E.** |
| haplo > 0.001 + SNPs | 0.67 | 0.063 | 0.65 | 0.082 |
| haplo > 0.005 + SNPs | 0.37 | 0.041 | 0.38 | 0.052 |
| haplo > 0.01 + SNPs | 0.30 | 0.029 | 0.31 | 0.037 |
| haplo > 0.05 + SNPs | 0.20 | 0.017 | 0.21 | 0.027 |
| SNPs Only | 0.20 | 0.019 | 0.20 | 0.024 |

To assess the possible impact of assay artifacts on our results we tested for the presence of spurious heritability (both *h*_*g*_^2^ and *h_hap_^2^*) between the two control cohorts (NBS and 58C; 2,635 controls and 2,794 controls, respectively). This estimate was not significantly different from zero (see Table 4), suggesting that control-specific assay artifacts are unlikely to contribute significantly to our estimates, although this analysis cannot rule out assay artifacts specific to disease cases. We sought to further ensure that the effects of assay artifacts were minimal by re-estimating *h*_*g*_^2^ and *h_hap_^2^* from sets of genotyped SNPs and haploSNPs obtained after excluding an additional 60k SNPs with even more stringent QC^9^ (see Online Methods). We compared these estimates to estimates obtained by dropping 10 random sets of 60k SNPs and observed no significant difference (see Table S5). This suggests that assay artifacts may not be contributing heavily to the observed increase in heritability explained, though data from other cohorts will be required to validate this result. As in our PGC2-SCZ analysis, the inclusion of rare haploSNPs did not decrease the contribution of common SNPs (see Table S6), again suggesting that heritability at common SNPs is not explained by rare variants

**Table 4.** Control-Control Heritability Estimates in WTCCC2 Controls

| **Multiple Sclerosis** |  |  |
| --- | --- | --- |
| **Model Components** | **$h_{\text{hap}}^2$** | **S.E.** |
| haplo > 0.001 + SNPs | -0.027 | 0.077 |
| haplo > 0.005 + SNPs | 0.019 | 0.051 |
| haplo > 0.01 + SNPs | -0.003 | 0.036 |
| haplo > 0.05 + SNPs | 0.004 | 0.028 |

We also estimated the heritability explained by genotyped SNPs and haploSNPs in a subset of ancestry matched cases and controls (see Online Methods). In this set of 8,149 individuals, we obtained *h*_*g*_^2^ estimates of *h*_*g*_^2^ = 0.20 (S.E. 0.024) and *h_hap_^2^* = 0.65 (S.E. 0.082) for haploSNPs with MAF > 0.001 (see Table 4), consistent with estimates obtained on the full data set. Thus, it is unlikely that our results are affected by the severe population stratification present in the overall set of 14,526 samples.

For further validation, we analyzed 38 loci (excluding chromosome 6, which contains the HLA locus) representing genome-wide significant associations from a larger sample^29^, as described above in the PGC2-SCZ analysis. The heritability explained by both SNPs and haploSNPs in these loci was estimated at *h h_ap_^2^* = 0.020 (S.E. 0.007), not significantly different than the heritability explained by SNPs alone *h*_*g*_^2^ = 0.029 (S.E. 0.004), again consistent with reports that the causal allele frequency spectrum at GWAS loci is skewed towards common variants^15^. We also investigated whether haploSNPs improved association signals at these loci by tagging unobserved causal variants. We compared whether the best haploSNP *χ*^2^ statistic was significantly larger than the best SNP *χ*^2^ statistic as described in the PGC2-SCZ analysis. Our results show that the improvement in *χ*^2^ statistics (mean improvement 3.6; mean increase of 32%) was statistically significantly different than expected due to the larger number of haploSNPs alone (K-S test P = 0.009; see Methods). As in the PGC2-SCZ analysis, this suggests that haploSNPs may tag causal variants better than SNPs alone.

Given the improvement in *χ*^2^ statistics at known associated loci, we sought to compare the effectiveness of association using haploSNPs vs. genotyped SNPs. We performed association of 375k SNPs and 38.8M haploSNPs (including chromosome 6) (see Figure 4). At a genome-wide significance threshold of 5e-8 for SNPs and 1.3e-9 for haploSNPs (0.05 / 38.8M) we observed only 4 loci containing genome-wide significant haploSNP associations compared with 7 loci containing genome-wide significant SNP associations. We note that this reduction in power is primarily due to correction for the larger number of hypotheses tested. To further explore the utility of haploSNP association, we relaxed our criteria and examined the set of SNPs and haploSNPs that achieved significance at an false discovery rate (FDR) 0.05^30^. At this FDR, we observed 82 SNP associations (see Table S7) and 260 haploSNP associations (see Table S8). Of the 260 haploSNP associations, 45 overlapped with known GWAS loci (see Online Methods), significantly more than the 28 SNP associations that did so (Permutation P = 6.15x10^−4^; see Online Methods). Additionally, the remaining 215 haploSNP associations (that did not overlap with known genome-wide significant loci) tended to overlap with larger numbers of genes than would be expected by chance, after accounting for the non-random distribution of SNPs and haploSNPs across the genome (Permutation P=0.009; see Online Methods). Textual analysis of all 260 loci using GRAIL^31^ implicated 19 genes with immune-related roles at 10 previously identified loci and implicated 5 immune related genes at 5 novel loci (see Table S9). We caution that an FDR of 0.05 does not meet traditional standards of genome-wide significance, and that these suggestive findings require replication in larger datasets; nonetheless, many of the novel loci are likely to be real, and these results suggest that haploSNPs may provide additional power to detect genetic associations to complex traits. While our modest sample size and the large multiple hypotheses correction penalty reduced our power to detect genome-wide significant associations, as sample sizes grow larger, the relative power of haploSNP association may increase.

**Figure 4.**
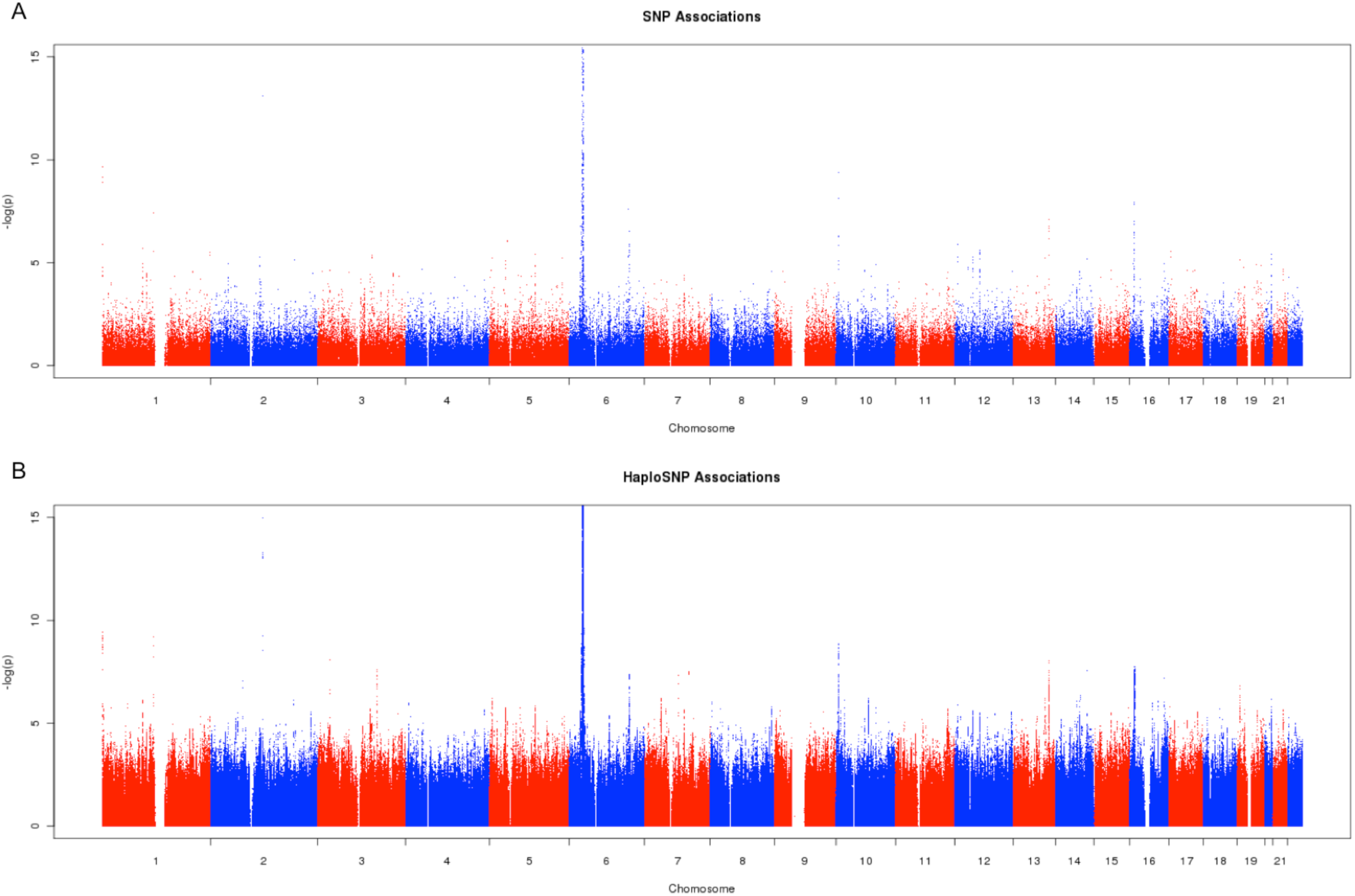
We compared the utility of haploSNPs association to that of SNPs alone in mapping loci for Multiple Sclerosis. At an FDR < 0.05, we observe 260 FDR controlled haploSNP associations, compared with 82 FDR controlled SNP associations suggesting that haploSNP association may provide additional power to discover genetic associations with complex traits.

## Discussion

We have developed a method for estimating the heritability explained by a specific set of haplotypes constructed from common SNPs (haploSNPs). Simulations and analyses of empirical schizophrenia and multiple sclerosis data sets show that these haploSNPs explain substantially more heritability than genotyped SNPs (or well-imputed SNPs). The absolute magnitude of increases in the heritability explained by haploSNPs for schizophrenia are similar to those that we observed in simulations with large numbers of rare variants; the observed increases in heritability explained for multiple sclerosis are even larger. One possible explanation for the large magnitude of these increases is that strong case-control ascertainment (especially for multiple sclerosis, whose prevalence is 0.1%) causes causal rare variants to have substantially higher frequency in the sample than in the population. We caution that, despite our efforts, assay artifacts may play a role in our multiple sclerosis estimates, and conclusive validation must be obtained via cross-cohort analyses of additional data sets, as we have provided here for schizophrenia.

Recent work has explored the alternative approach of including low-accuracy imputed SNPs in estimates of heritability explained by SNPs, and has shown that this approach can explain substantial additional heritability for quantitative traits^32^. However, it is unclear whether this approach can work for case-control traits due to the severe danger of assay artifacts inflating the resulting estimates^9^. Additionally, imputation will not capture the contributions of variants that appear rarely in, or are entirely absent from reference panels. This is particularly relevant for large-effect rare variants for diseases of low prevalence that will be at dramatically higher frequency in ascertained case-control samples than in the entire population. In these instances, haploSNPs may be a more effective approach to tagging unobserved causal variants.

We note that explaining more heritability using a specific set of haploSNPs (*h*_*hap*_^2^) is distinct from estimating total narrow-sense heritability (*h*^2^), and that estimates of *h*_*hap*_^2^ should not be viewed as unbiased estimates of *h*^2^. While haploSNPs may not capture all of trait heritability, specific haploSNP associations should explain all of the estimated haploSNP heritability in the limit of infinite sample size. Additionally, unbiased estimates of *h*^2^ are typically based on long haplotypes shared identical-by-descent (IBD) between closely related individuals^11,33^ and may be subject to biases due to shared environment^11^ or genetic interactions that are difficult to tease out^10^. By contrast, approaches based on unrelated individuals, such as ours, are unaffected by these biases. We view the use of IBD analyses to produce estimates of *h*^2^ in unrelated individuals as an exciting avenue for future work.

Although haploSNPs explain substantially more heritability than genotyped SNPs, this approach does have limitations. First, as with other heritability analyses of case-control traits, the approach can potentially be susceptible to assay artifacts, motivating cross-cohort analyses such as we have performed here with schizophrenia. Second, haploSNPs could in theory be tagging effects due to interaction between co-located SNPs, though the bulk of these effects may still be likely to be largely additive^34^. Third, efforts to leverage haploSNPs to detect new associations will suffer a substantial multiple hypothesis testing burden due to the large number of haploSNPs, potentially requiring large sample sizes to overcome; however, the 215 suggestive (FDR< 0.05) novel loci that we detected in a multiple sclerosis data set of modest sample size suggest that overcoming this multiple hypothesis testing burden may be feasible. Fourth, construction of haploSNPs from well-imputed SNPs (as would be necessary for haploSNP association meta-analysis of cohorts typed on different genotyping arrays) remains a direction for future research; for this reason, in our PGC2-SCZ analysis we meta-analyzed *h*_*hap*_^2^ estimates without attempting a haploSNP association meta-analysis. Despite these limitations, we anticipate that haploSNPs will be a valuable approach for rare variant analysis as increasingly large genotyping array data sets (e.g. UK Biobank, 500k samples; see Web Resources) are generated as a precursor to large-scale sequencing studies.

## Online Methods

### Definition of haploSNPs

haploSNPs are haplotypes of adjacent SNPs excluding a subset of masked sites that arise from skipped mismatches. Individuals are considered to carry 0,1, or 2 copies of the haploSNP if none, one or both of their chromosomes matches the haplotype at all unmasked sites.

### Algorithm to generate haploSNPs

The algorithm to generate haploSNPs (see Supplementary Material for pseudocode) proceeds from phased genotypes. For our haploSNP analysis, we used the HAPI-UR method^17^ to computationally phase genotypes. Using these phased genotypes we compare every pair of chromosomes searching for shared segments. A shared segment is begun at a single SNP at which the two chromosomes match alleles. This segment is extended in one direction until a terminating mismatch between the chromosomes is found. A terminating mismatch is one that cannot be explained without a recombination between the current segment and the mismatch SNP. This is tested using a standard 4-gamete test^35^. That is, for all chromosomes in the sample (not just the pair under consideration) we assign a 1 or 0 allele for the segment being extended (*current_haplo* in pseudocode). A 1 allele is assigned to chromosomes that perfectly match the segment at all sites and a 0 is assigned to all other chromosomes. We compare this set of alleles to those at the mismatch SNP. Given two alleles at the haploSNPs and two at the mismatch SNP, a maximum of four possible allelic combinations can be observed. If all four combinations are observed, this indicates that a recombination event is required to explain the mismatch, and the haploSNP will be terminated. If, however, only three combinations are observed, the mismatch may be explained by a mutation on the shared haplotype background. These mismatches are ignored and the haploSNP is extended further. We note that this approach can produce a very large number of haploSNPs and very long haploSNPs that could tag signals of cryptic relatedness. To reduce the number of haploSNPs, making analysis tractable, we used a max length of 50kb. We note that haploSNPs that are truncations of prior haploSNPs—starting at a later SNP but matching the exact same set of individuals over the same subsequent set of SNPs—are excluded from output.

The output of this algorithm is a list of haploSNPs. These haploSNPs are made up of multiple co-located SNPs and a particular allele at each SNP. For each haploSNP, each phased chromosome is assigned either a 1, indicating a perfect match at all SNPs that make up the haploSNP, or a 0 otherwise. This set of biallelic haploSNPs is then used in downstream analysis in addition to biallelic SNPs. We note that all quality control steps (see below) are applied to SNPs prior to construction of haploSNPs, no additional QC steps are applied to the haploSNPs in our analysis.

### Linear mixed model for estimating components of heritability

We summarize the model the generative model for quantitative traits that is common to PCGC regression^18^ and REML estimation^7,25^. Formally, we assume that the phenotype is generated from a model *y* = Σ*_i_ β_i_W_i_* + *e* where *β_i_* is the effect size *W*_*i*_ is the standardized genotype for SNP *i* and *e* is environmental noise. We can then describe the phenotypic variance-covariance matrix 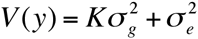 where *K* represents the exact sample covariance over causal variants, 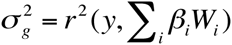, and the narrow-sense heritability is 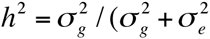). For case-control traits, we assume a liability threshold model^9^ where the underlying liability is a quantitative trait as described above and cases are defined as those individuals whose liability exceeds a pre-defined threshold. Because of bias in REML estimation introduced by case-control ascertainment, we used PCGC regression to produce estimates^18^.

### MAF-stratified PCGC regression

Separately from the generative model, we can consider the estimation procedure where we seek to estimate the heritability explained by a specific set of variants, *s*. In this case, 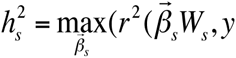), which is the maximum phenotypic variance explained by a linear combination of variants in *s*. Invoking this linear model, we write the phenotypic variance-covariance as 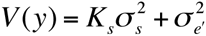 where 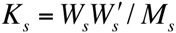 is the genetic kinship matrix calculated for this specific set of variants, *M*_*s*_ is the number of variants in the set, 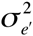 represents all phenotypic variance not captured by variants in *s*, and *y* is normalized to have mean 0 and variance 1. We note that 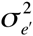 may be environmental noise or genetic variance due to causal variants not well captured by linear combinations of variants in *s*. PCGC regression uses this equation for the phenotypic variance-covariance matrix, regressing *V* (*y*) against *K*_*s*_. The estimated slope in this regression is an estimate of 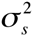 To partition the genetic variance between categories of variants, we simply extend this to perform a multiple regression of 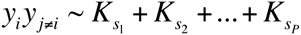 obtaining estimates of the variance explained by each partition 1, 2,…, *P*, which are given by the regression coefficients. Note that in this regression, only the off-diagonal entries of the phenotypic and genetic covariance matrices are used.

We stratified our haploSNPs by MAF to avoid LD-bias due to MAF-dependent genetic architectures^21,22^. When analyzing PGC2-SCZ and WTCCC2-MS data sets, we considered MAF bins of 0.001-0.005, 0.005-0.01, 0.01-0.05, 0.05-0.1, 0.1-0.25, and 0.25-0.5. When analyzing the UK10K data-set we also included a bin for MAF < 0.001. For each MAF bin we computed a separate kinship matrix, (i.e. 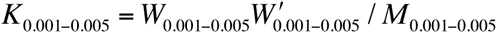). And we fit a joint model for all MAF bins, estimating using multiple regression (see above). Standard errors for PCGC regression were estimated using a jackknife procedure over individuals^18^. This procedure produces estimates of the standard error for each component as well as the sum across all components.

REML estimation uses the same model as described above, but assumes a multivariate normal distribution for the phenotype, genotypic effects and environmental effects, 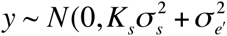). Then the variance component 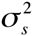 can be estimated by maximizing the likelihood of the data according to this distribution. While this method can produce more precise estimates if the underlying normality assumption holds, this assumption is often violated when analyzing ascertained case-control traits^18^. As we observed substantial biases in heritability estimates using REML, we elected to use MAF-stratified PCGC regression for analyses described in the main text. However, estimates were also computed using MAF-stratified REML and are presented in Tables S1 and S2.

### Assessing tagging in simulations

We sought to quantify the improvements in tagging of rare variants when using haploSNPs as compared with genotyped SNPs alone by analyzing a large sample of sequenced individuals from the UK10K project.

We began with 235k SNPs on chromosome 22 genotyped in 3047 unrelated individuals. We considered three possible sets of tags:

1. 6477 SNPs that were present on the Illumina Human660-Quad chip
2. 24k well imputed SNPs. Imputation was performed using a separate phasing with SHAPEIT^36^ followed by imputation with IMPUTE2^23^. SNPs with INFO < 0.9 were excluded as were SNPs that had P < 0.01 for deviation from Hardy-Weinberg equilibrium. No allele frequency threshold was imposed.
3. 937k haploSNPs that were built from the SNPs in (1). Specifically, these “genotyped SNPs” were computationally phased^17^ and then used to construct haploSNPs with length < 50kb according the algorithm given above.

For each of the 235k SNPs in the sequence data, we assessed the best tag in 1Mb window surrounding the SNP using each set of tags. When using haploSNPs we observed a highly statistically significant increase in the quality of the best tag (see Figure 1). This was particularly pronounced for rare and low-frequency variants.

To ensure that observed improvements in tagging of rare variants were not due to chance, we ran permutation experiments with the haploSNPs. Specifically, we permuted haploSNPs but left SNPs intact and re-assessed the average best-tag for each of the 235k sequence SNPs. This produced a set of random haploSNPs of the same size and with the same allele frequency spectrum of the original haploSNPs. Over 5 permutations, we observed an average best tag of *r^2^* =0.37 (s.d. = 0.0002), suggesting that the observed improvement (*r^2^* =0.54) is not driven primarily by improved tagging by chance.

### Robustness simulations

We sought to ensure that PCGC regression produced robust estimates of the heritability explained by haploSNP under a variety of different genetic architectures. We used the same 937k haploSNPs generated above. These haploSNPs were then treated as though they were, in fact, genotyped SNPs and phenotypes were simulated using these haploSNPs as causal variants. Thus, the proportion of phenotypic variance explained by haploSNPs was known exactly and could be compared to the estimated *h_hap_^2^*. For each set of 100 simulated phenotypes, we chose a range of the frequency spectrum to draw causal haploSNPs from. Frequency ranges were MAF > 0.001, 0.01 and MAF < 0.05, 0.1, and 0.5. Once causal variants were selected, effect sizes were assigned using a normal distribution:

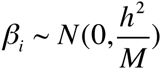

where *h*^2^ is the heritability of the simulated phenotype, *M* is the number of causal haploSNPs used, and *β*_*i*_ is the effect size for genotypes after standardization to mean 0 and unit variance. This ensures that causal SNPs explain equal variance, on average, regardless of MAF. For these simulations we used values of *h*^2^ = 0.5 and *M* = 1000. For each simulated phenotypes, we fit a model with six MAF stratified variance components as described above. The resulting heritability estimates were summed across all components.

### Assessing increases in explained heritability in simulation

To assess our ability to explain additional heritability using haploSNPs, we simulated phenotypes from UK10K sequence SNPs (instead of haploSNPs as above). We simulated phenotypes using a variety of genetic architectures with different levels of rare variant involvement. For each set of 100 simulated phenotypes, we chose a range of the frequency spectrum to draw causal SNPs from. Frequency ranges were MAF > 0.001, 0.01, 0.05 and MAF < 0.05, 0.1, 0.2, 0.3, 0.4, and 0.5, as above. Within these ranges, variants were selected at random from the underlying SNPs in sequence. Once causal variants were selected, effect sizes were assigned as described above. For these simulations we used values of *h* ^2^ = 0.5 and 1000 causal variants.

Once phenotypes were generated, we examined a set of 6477 genotyped SNPs and a set of 932k haploSNPs (see above). We estimated the heritability explained by SNPs alone, or haploSNPs + SNPs using MAF-stratified PCGC regression^18^.

We also explored the effects of varying the length threshold on the performance of haploSNPs in tagging additional heritability. Our results show that while longer haploSNPs can tag more heritability, they do so at the cost of a substantial increase in standard error (see Figure S2). Based on this, all of our primary analyses (simulations + real data) restricted to haploSNPs of length < 50kb. We note that this choice was a function of sample size, and future studies at larger sample sizes would be able to include longer haploSNPs while keeping standard errors manageable.

Finally, we note simulation of ascertained case-control traits is challenging. To perform such a simulation with severe case-ascertainment using real genotypes we would need a data-set with large numbers of individuals with observed rare variants. Such a data-sets are not currently available.

### Data sets

For simulations we analyzed data from the UK10K project. For real data analyses, we examined data from the Psychiatric Genomics Consortium 2 (PGC2) and the Wellcome Trust Case Control Consortium 2 (WTCCC2). These data have been previously described in ^3^ and ^29^. Because heritability analyses can be particularly susceptible to artifacts, we applied additional quality control (see below) to each data set.

#### PGC2 Large Cohort Meta-analysis Data

The cohorts comprising the PGC2 have diverse European ancestry and were genotyped on a variety of different platforms. To avoid issues related to cross-population heritability estimation, we focused on meta-analysis of estimates within cohorts. Our estimates were produced from each of 10 cohorts with >1,000 individuals.

Within each cohort we applied stringent QC to SNPs, removing any SNPs that were below 0.01 minor allele frequency, above 0.002 missingness, had deviation from Hardy-Weinberg equilibrium at a p-value below 0.01, or had differential missingness between cases and controls with a p-value below 0.05. We also removed one individual in any pair of individuals with relatedness greater than 0.025 by SNP covariance. Following all QC steps, we analyzed 35,238 individuals. The average number of SNPs genotyped in each cohort was 461k. We did not use imputed SNPs to build haploSNPs, as we were concerned that it would be circular to use genotypes inferred from haplotype structure as a basis to infer haplotype structure.

#### PGC2 Cross-cohort Analysis Data

Our cross-cohort analysis was designed to merge PGC2 cohorts of homogenous ancestry that had significant overlap in their sets of genotyped SNPs. We chose 2 subsets of the PGC2 data. The first subset was made up of five cohorts of Swedish ancestry and consisted of five cohorts with a total of 2819 schizophrenia cases and 2911 controls. The second subset was of British ancestry and consisted of two cohorts with a total of 5406 treatment resistant schizophrenia cases and 5284 controls.

Within each subset we applied stringent QC to SNPs as described above: removing any SNPs that were below 0.01 minor allele frequency, above 0.002 missingness, had deviation from Hardy-Weinberg equilibrium at a p-value below 0.01, or had differential missingness between cases and controls with a p-value below 0.05. We also removed any pair of individuals with relatedness greater than 0.025 by SNP covariance. Following QC, we analyzed genotypes at 310k and 303k SNPs in each subset, respectively.

#### WTCCC2 Quality Control

Because data from additional cohorts was not available for validation of heritability estimates, we used a higher level of stringency than described above. Specifically, we removed any SNPs that had minor allele frequency below 0.02, had missingness greater than than 0.002, had deviation from Hardy-Weinberg equilibrium at a p-value below 0.05, or had differential missingness between cases and controls with a p-value below 0.05. These thresholds for SNP QC were decided upon after analyzing 3 levels of increasingly stringent QC. We chose this level of QC because removing SNPs in further rounds of increasingly stringent QC did not alter results compared with removing random SNPs (see Table S4). To deal with subtle relatedness, we removed one individual of any pair that showed relatedness > 0.05 as estimated by SNP covariance. This was different than above We subsequently performed five rounds of outlier removal whereby all individuals more than 6 standard deviations away from the mean along any of the top 20 eigenvectors were removed and all eigenvectors recomputed. Following all QC steps, we analyzed 14,526 individuals genotyped at 375k SNPs. These individuals consisted of 9315 Cases, 2635 controls from the NBS cohort and 2794 controls from the 58C cohort. To avoid the effects of the well-known HLA locus, we excluded chromosome 6 from all heritability analyses, leaving a total of 349k SNPs. While the effect of the HLA locus is the largest in the genome, it has been estimated to explain only about 3% of the phenotypic variance of MS on the liability scale^37^, and thus should not affect our results substantially.

### Accounting for population stratification in real data

To prevent population stratification from influencing our estimates of heritability for PGC2-SCZ and WTCCC2-MS we computed the top 20 principal components (PCs) for the kinship matrix computed from each of the MAF bins of haploSNPs described above and the top 20 PCs for a kinship matrix computed from genotyped SNPs. This total of 140 PCs was used to adjust the PCGC regression using a logistic model of fixed effects in an ascertained study as described in Golan et al. 2014 PNAS^18^. In addition, as recommended by these authors, we subtracted these top 20 PCs from each kinship matrix weighted by eigenvalue. That is, we replaced a kinship matrix for a set of variants *s* with 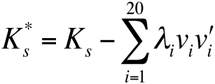. We note that this is akin to fitting each SNP as a linear combination of the top 20 PCs and including the residuals to compute the kinship.

### Assessing cross cohort heritability

To ensure that observed increases in heritability explained in schizophrenia were due to tagging of causal variation and not cohort-specific assay artifacts, we estimated heritability across cohorts within homogenous ancestry subsets. Specifically, we used a modified version of the standard MAF-stratified PCGC regression technique to perform regression. In the standard regression,

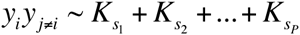

off-diagonal entries of the phenotypic covariance matrix are regressed against off-diagonal entries of the genetic covariance matrix. To assess cross-cohort heritability, we performed the identical regression, simply using all pairs of individuals drawn from a different original cohort.

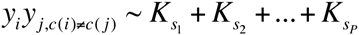

where *c(i)*, *c(j)* are the cohorts to which individuals *i,j* belong.

### Analysis of GWAS Loci

We sought to assess whether haploSNPs at known associated loci explained significantly more heritability than SNPs alone. Given genome-wide significant loci extracted from recent analyses^3,29^, we used regions of ±500kb from the GWAS SNP. Overlapping regions were merged. These SNPs were then used to construct haploSNPs. These haploSNPs were stratified into MAF bins (see above) and heritability was estimated using PCGC regression. Two models were tested: (1) a 7 variance component model with 6 MAF stratified haploSNP components and 1 SNP component and (2) a 1 variance component model with 1 SNP component. Principal components analysis was performed on these GRMs and the top 20 PCs were subtracted from the genetic relationship matrices, after weighting by eigenvalue (see above). For schizophrenia, heritability was estimated across cohorts (see above).

To assess whether haploSNPs better tagged causal variants at GWAS loci, we performed association using logistic regression after correcting for principal components and compared the best haploSNP *χ*^2^ statistic to the best SNP *χ*^2^. To assess the significance of the observed improvement, we simulated phenotypes using the SNP with the highest SNP *χ*^2^ as the only causal variant. The effect size of the SNP was chosen to match the observed effect size. For each simulated phenotype, we performed association and assessed the best haploSNP *χ*^2^ and the best SNP *χ*^2^. We performed simulations until at least 100 simulations showed larger improvements than observed in real data, or until 10,000 simulations were performed. The significance was then assessed as:

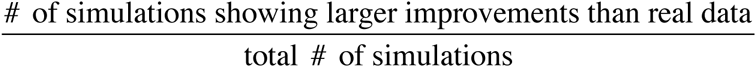

These p-values were meta-analyzed across cohorts and loci and compared with the uniform distribution using the Kolmogorov-Smirnov test (alternative “greater” for the CDF to test whether P-values are systematically smaller than expected).

### haploSNPs Association Analyses

haploSNP and SNP associations were performed using logistic regression implemented in the PLINK2 software package (see Web Resources). 20 Principal Components inferred from SNPs were used to correct for population stratification. To control the false discovery rate, we used the Benajamini-Hochberg correction procedure^30^.

To assess the degree to which FDR controlled haploSNP associations increased the overlap with previously published loci, we permuted the location of haploSNP associations. Specifically, the 199 haploSNP associations that did not overlap with SNP associations were replaced with 199 random haploSNPs. No pair of random haploSNPs was chosen to be closer than 500kb. For each random assignment, the overlap with known loci was assessed. An association was considered overlapping if the middle of the haploSNP was within 500kb of the association. The associations were the combined set of loci published in Sawcer et al. 2011 *Nature* ^29^ and the MS loci analyzed in Gusev et al. 2013 *PLoS Gene* t^15^. The p-value was estimated as the proportion of random assignments showing a larger increase than observed in real data.

We also sought to assess the degree to which FDR controlled haploSNP associations showed genic enrichment. For each of the 215 haploSNP associations that did not overlap with known loci, we assessed the number of genes within a 1MB window (500kb on either side). We compared this to the number of genes within a 1MB window of 215 randomly selected haploSNPs.

## Web Resources

HaploSNP Software: www.hsph.harvard.edu/faculty/alkes-price/software/

Efficient PCGC Regression Software: http://www.hsph.harvard.edu/faculty/alkes-price/software/

UK10K data set: http://www.uk10k.org

UK Biobank:http://www.ukbiobank.ac.uk

PLINK2: www.cog-genomics.org/plink2

## References

1. Visscher, P. M., Brown, M. A., McCarthy, M. I. & Yang, J. Five years of GWAS discovery. Am. J. Hum. Genet. 90, 7–24 (2012).

2. Sullivan, P. F., Kendler, K. S. & Neale, M. C. Schizophrenia as a complex trait: evidence from a meta-analysis of twin studies. Arch. Gen. Psychiatry 60, 1187–1192 (2003).

3. Consortium, S. W. G. O. T. P. G. Biological insights from 108 schizophrenia-associated genetic loci. Nature 511, 421–427 (2014).

4. Polderman, T. J. C. et al. Meta-analysis of the heritability of human traits based on fifty years of twin studies. Nature genetics 47, 702–709 (2015).

5. Maher, B. Personal genomes: The case of the missing heritability. Nature 456, 18– 21 (2008).

6. Manolio, T. A. et al. Finding the missing heritability of complex diseases. Nature 461, 747–753 (2009).

7. Yang, J. et al. Common SNPs explain a large proportion of the heritability for human height. Nature genetics 42, 565–569 (2010).

8. Lee, S. H. et al. Estimating the proportion of variation in susceptibility to schizophrenia captured by common SNPs. Nat Genet 44, 247–250 (2012).

9. Lee, S. H., Wray, N. R., Goddard, M. E. & Visscher, P. M. Estimating missing heritability for disease from genome-wide association studies. Am. J. Hum. Genet. 88, 294–305 (2011).

10. Zuk, O., Hechter, E., Sunyaev, S. R. & Lander, E. S. The mystery of missing heritability: Genetic interactions create phantom heritability. Proceedings of the National Academy of Sciences (2012).

11. Zaitlen, N. et al. Using extended genealogy to estimate components of heritability for 23 quantitative and dichotomous traits. PLoS Genet 9, e1003520 (2013).

12. Zaitlen, N. et al. Leveraging population admixture to characterize the heritability of complex traits. Nature genetics 46, 1356–1362 (2014).

13. Gibson, G. Rare and common variants: twenty arguments. Nature reviews. Genetics 13, 135–145 (2012).

14. Marchini, J. & Howie, B. Genotype imputation for genome-wide association studies. Nature reviews. Genetics 11, 499–511 (2010).

15. Gusev, A. et al. Quantifying Missing Heritability at Known GWAS Loci. PLoS Genet 9, e1003993 (2013).

16. Powell, J. E., Visscher, P. M. & Goddard, M. E. Reconciling the analysis of IBD and IBS in complex trait studies. Nature reviews. Genetics 11, 800–805 (2010).

17. Williams, A. L., Patterson, N. & Glessner, J. Phasing of Many Thousands of Genotyped Samples. The American Journal of … (2012).

18. Golan, D., Lander, E. S. & Rosset, S. Measuring missing heritability: Inferring the contribution of common variants. Proceedings of the National Academy of Sciences 111, E5272–81 (2014).

19. Haseman, J. K. & Elston, R. C. The investigation of linkage between a quantitative trait and a marker locus. Behavior Genetics 2, 3–19 (1972).

20. Yang, J., Zaitlen, N. A., Goddard, M. E., Visscher, P. M. & Price, A. L. Advantages and pitfalls in the application of mixed-model association methods. Nature genetics 46, 100–106 (2014).

21. Speed, D., Hemani, G., Johnson, M. R. & Balding, D. J. Improved Heritability Estimation from Genome-wide SNPs. The American Journal of Human Genetics 91, 1011–1021 (2012).

22. Lee, S. H. et al. Estimation of SNP heritability from dense genotype data. Am. J. Hum. Genet. 93, 1151–1155 (2013).

23. Howie, B. N., Donnelly, P. & Marchini, J. A flexible and accurate genotype imputation method for the next generation of genome-wide association studies. PLoS Genet 5, e1000529 (2009).

24. 1000 Genomes Project Consortium et al. An integrated map of genetic variation from 1,092 human genomes. Nature 491, 56–65 (2012).

25. Yang, J., Lee, S. H., Goddard, M. E. & Visscher, P. M. GCTA: a tool for genome-wide complex trait analysis. Am. J. Hum. Genet. 88, 76–82 (2011).

26. Loh, P. R., Bhatia, G., Gusev, A. & Finucane, H. K. Contrasting regional architectures of schizophrenia and other complex diseases using fast variance components analysis. bioRxiv (2015).

27. Clayton, D. G. et al. Population structure, differential bias and genomic control in a large-scale, case-control association study. Nat Genet 37, 1243–1246 (2005).

28. Lee, S. H., Yang, J., Goddard, M. E., Visscher, P. M. & Wray, N. R. Estimation of pleiotropy between complex diseases using single-nucleotide polymorphism-derived genomic relationships and restricted maximum likelihood. Bioinformatics 28, 2540–2542 (2012).

29. International Multiple Sclerosis Genetics Consortium et al. Genetic risk and a primary role for cell-mediated immune mechanisms in multiple sclerosis. Nature 476, 214–219 (2011).

30. Benjamini, Y. & Hochberg, Y. Controlling the False Discovery Rate: A Practical and Powerful Approach to Multiple Testing on JSTOR. Journal of the Royal Statistical Society Series B … (1995).

31. Raychaudhuri, S., Plenge, R. M., Rossin, E. J. & Ng, A. C. PLoS Genetics: Identifying Relationships among Genomic Disease Regions: Predicting Genes at Pathogenic SNP Associations and Rare Deletions. PLoS Genet (2009).

32. Yang, J. et al. Estimation of genetic variance from imputed sequence variants reveals negligible missing heritability for human height and body mass index. Nature Genetics, submitted

33. Price, A. L., Helgason, A., Thorleifsson, G. & McCarroll, S. A. PLoS Genetics: Single-Tissue and Cross-Tissue Heritability of Gene Expression Via Identity-by-Descent in Related or Unrelated Individuals. PLoS genetics (2011).

34. Hill, W. G., Goddard, M. E. & Visscher, P. M. Data and theory point to mainly additive genetic variance for complex traits. PLoS Genet 4, e1000008 (2008).

35. Richard R Hudson, N. L. K. Statistical Properties of the Number of Recombination Events in the History of a Sample of DNA Sequences. Genetics 111, 147 (1985).

36. Delaneau, O., Marchini, J. & Zagury, J.-F. A linear complexity phasing method for thousands of genomes. Nat. Methods 9, 179–181 (2012).

37. Lee, S. H. et al. Estimation and partitioning of polygenic variation captured by common SNPs for Alzheimer’s disease, multiple sclerosis and endometriosis. Human molecular genetics 22, 832–841 (2013).

